# A Community-based assessment of quality of health care received and client satisfaction in Lagos, Nigeria

**DOI:** 10.1101/541565

**Authors:** Modupe Rebekah Akinyinka, Esther Oluwakemi Oluwole, Olumuyiwa Omotola Odusanya

## Abstract

**Background:** Quality of health care and client satisfaction are key elements in improving the performance of health systems. A community-based assessment was conducted to determine the level of client satisfaction and the perception of the quality of services received by citizens of Lagos State.

**Methodology:** A descriptive cross-sectional study using both quantitative and qualitative methods, was conducted in four local government areas of Lagos state selected by multi-stage sampling techniques. The survey instruments included an interviewer- administered, pre-tested questionnaire and a 10-itemed focus group discussion guide. The survey obtained information about quality of the facilities and services. The outcome variables were client satisfaction and service quality.

**Results:** Two thousand respondents were recruited with a mean age of 37.6±10.21 years. Almost all respondents (98%) rated the health facilities to be clean, 96% felt they received effective treatment from their providers. Six out of ten respondents rated the waiting time to be short and 60% felt that most drugs were available. Eight-five percent opined that the quality of care received was good and 95% were satisfied with the services received. There was a significant correlation between quality of care and client satisfaction (r=0.145, p=0.001). Service factors such as perceived effectiveness of treatment received, confidence in health providers and use of higher level of health care were predictive of client satisfaction and good service quality.

**Conclusion:** Most clients were satisfied with health services and felt that service quality was good.

## Introduction

The quality of a health system reflects the values and goals current in the medical care system and in the larger society of which it is a part. In addition, quality is an evaluation of the gap between service expectation and performance. As quality does not readily lend itself to an easy definition, assessment of quality must rest on a conceptual and operationalized definitions of what the quality of medical care means.^1^

A survey amongst 239 users of health services in the United States of America (USA) on the meaning of quality health care revealed diverse opinions such as access to health care, having competent and skilled providers and receiving proper treatment showing that perception of quality varies and revolves around competence and skills of providers.^2^ Another researcher identified communication with patients, competence of staff, demeanor of staff, quality of the facilities and perceived costs as significant factors that explain variation in customer satisfaction with hospitals.^3^

Taylor posits that service quality and customer satisfaction should be viewed as separate constructs. He stated further that service quality perceptions should be considered as long-term consumer attitudes, whereas patient satisfaction deals with short-term, service-encounter-specific consumer judgments.^4^ While this may be true, there is no one way to correctly separate the two elements as they are interwoven particularly for the acutely ill patient. Woolley et al noted that patient satisfaction was a product of four variables: satisfaction and outcome, continuity of care, patient expectations, and doctor-patient communications.^5^

Mosadeghrad developed a framework for measurement of quality in health care. The framework includes both tangibles and intangible elements. The intangible elements were the environment, empathy, efficiency, effectiveness and courtesy.^6^ The intangible elements are often not easy to measure and require a lot of observations to make unbiased decisions.

Furthermore, they do not act independently of each other but are moderated by other factors as reported by Tucker and Adam, who observed that other moderating factors apart from patients’ socio-demographic characteristics could predict client satisfaction and quality.^7^ Furthermore, provider performance and access to health care were found to capture 74% of the variance in satisfaction quality and were positively associated with patients’ assessment of satisfaction of quality (p=0.001). Patients’ socio-demographic differences were found to account for only 1% of the variance.^7^

Parasuvaman et al developed five generic domains of service quality (SERVIQUAL) namely tangibles, reliability, responsiveness, assurance and empathy.^8^ However, Bowers et al reported that although the elements of the generic SERVIQUAL dimensions are found in health care they do not completely define the constructs of health care quality. They identified empathy, reliability, responsiveness (in the SERVIQUAL model) along with communication and caring as five indicators of health care quality on a global satisfaction measure. ^9^ The client-focussed definition of quality comes from Donabedian et al whose model provides a framework for examining health services and evaluating quality of care. In this model, quality of care can be assessed in three major areas “structure,” “process,” and “outcomes”.^1, 10^

The quality and cost of health services are determinants of utilization of those services depending on the population surveyed. A survey amongst 840 households across selected urban, peri urban and rural communities, in southeast Nigeria, found that of the nine (9) demographic variables, only the locality and status of the health system (strong or weak in terms of child immunization) was found to influence both the poor rating and utilization of public health services. Individuals from states with strong health system rated relatively higher and used public health services more (p <0.001), than their counterparts from states with weak health care systems. Similarly, those in the urban or peri-urban localities used public health services more (p = 0.013). Clients with a good perception of the quality of health service provided, rated and patronized them more (p < 0.001).^11^

A study of waiting time and service satisfaction at antenatal clinics in Ile-Ife reported that only 55% of the women were satisfied with the quality of antenatal care received. There was prolonged waiting and transit times for the clients. About 72% of the women felt the service was good, 53% assessed staff to be competent and 39% felt that the staff were friendly. A higher level of education was found to be significantly (p =0.02) associated with satisfaction.^12^ A study of client perception of service quality at outpatient clinics of a general hospital in Lagos reported that 88% rated the overall service quality to be good with the assurance domain as the most important predictor (p <0.001) of service quality.^13^ Amole et al in their study of patients perception of service quality in six teaching hospitals in southwest Nigeria reported that the most important factor to patients was the reliability dimension, followed by the assurance dimension with the empathy dimension being the least important factor.^14^ This contrasts with another study by Oyatoye et al who studied four general hospitals in Ogun Sate and found that patients had the greatest preference for empathy while the least preference was waiting time.^15^

At the primary health care level, a study in north-central Nigeria reported that the highest perception was for lack of interruption during consultation while the lowest was in the domain of respect for persons. Seven out of ten patients felt that the consulting room provided enough privacy and 84% felt that they received adequate information from their doctors. Age, sex, educational level and income were significantly associated with satisfaction.^16^ A study conducted at a flagship primary health centre in urban Lagos found waiting time to be as long 138 minutes and was the least liked aspect of care provided in the facility by the larger proportion of the clients (32.9%).^17^

A few studies on quality of health services exist in Nigeria as previously highlighted, but many were limited to one health facility and conducted amongst clients of the facility studied. Therefore, this study was conceived as a community- based assessment of the perception of the quality of services received by citizens of Lagos State. It is hoped that the findings of this study will be useful for hospital management authorities, health planners and policy makers to improve patient experiences and the quality of health care provided.

## Materials and methods

### Background information to study area

Lagos State was created on May 27, 1967. It is in the Southwest geopolitical zone of Nigeria. It was the capital of Nigeria until 1991. Ikeja is the capital city of the State. Lagos remains the economic capital of Nigeria. The State has 20 Local Government Areas (LGA). Sixteen of the LGAs are classified as urban and four are rural. Health facilities are provided through a mix of private and public facilities at primary, secondary and tertiary levels.

### Study Design

The study design was a descriptive cross-sectional using both quantitative and qualitative methods to investigate client satisfaction, service characteristics, and the perception of health care quality received by community members in Lagos State. An interviewer-administered questionnaire was used to obtain information for the quantitative aspect of the study. Focus group discussions (FGDs) were held for the qualitative aspect.

### Study population and eligibility criteria

The study population was drawn from adult residents aged 18 years and above which were living in the selected LGAs.

#### Inclusion criteria

Consenting adults aged 18 years and above living in the selected LGAs.

#### Exclusion criteria

Residents who did not give consent.

### Sample size determination

The minimum sample size for quantitative data collection was determined using the appropriate formula for prevalence studies.^18^ The statistical assumptions for determining the minimum sample size were: a type 1 error rate of 5%, a prevalence of 0.58 of positive perception of health workers by community members,^19^ a precision of ±2.5 percentage points and a 20% non-response rate. Thus, the calculated minimum sample size was 1919, which was rounded up to 2000. The participants for the FGD were purposively selected. One FGD session was held in each LGA and the number of participants was averagely ten (10).

### Sampling techniques

A multi stage sampling method was used to select the subjects for quantitative data collection in this study. In the first stage, out of the 20 Local Government areas (LGAs) in Lagos state, of which 16 are urban and 4 are rural, four LGAs (three urban and one rural) were selected using stratified random sampling by balloting. These were Ikeja, Mushin, Ojo (urban) and Badagry (rural) LGAs. In the second stage, at each of the selected LGA, two wards were selected by simple random sampling by balloting. In the third stage, using the sampling frame of all streets in the selected wards, a minimum of 10 streets were selected by using a table of random numbers.

The fourth stage involved selecting consecutive houses on each street using the Local Government house numbering system starting from the first number. In the fifth stage, one household was selected by balloting and a consenting adult was approached to participate in the study. Where there was more than one consenting adult in the selected household, one was chosen by balloting. Twenty-five respondents were selected from each street, and an equal number (500) of respondents were selected per LGA to allow for equal representation from all selected areas. For qualitative data collection, one focus group discussion was held per LGA. FGDs were held for female participants in Mushin, Ojo and Badagry and for male participants in Ikeja. Ten participants were selected via purposive sampling based on willingness and availability to participate in each FGD session.

### Survey Instruments

Two instruments were developed for the study. The first was an interviewer- administered, pre-tested questionnaire and the second was a ten-itemed focus group discussion guide. The interviewer- administered questionnaire instrument was developed from a review of the literature on the subject and was pre-tested in Alimosho LGA. The alpha Cronbach reliability coefficient was 0.792. The instrument was modified and administered after pre-testing. The instrument had two sections. The first dealt with socio-demographic characteristics of the respondents such as age, gender, educational level and occupation. The second focused on utilization of health facilities, accessibility, preferred places for treatment of common health conditions, assessment of quality of the facilities and providers. Additional information was sought on client satisfaction and the quality of the service received. The FGD guide sought for information on the utilization of health facilities, facility environment (toilets, waiting areas, consulting rooms), competence and attitude of health workers, ease of using the facility and problems encountered by respondents during visits to health facilities.

### Data collection

The quantitative data was collected by four trained research assistants (who had a minimum of secondary school education) between February and March 2017. Research assistants were trained for two days prior to data collection. Participants for the FGD were invited and reminded via text messages and calls. The selected participants were within the same age range for each FGD. All sessions were audio recorded after obtaining written informed consent from the participants.

### Data management

All completed questionnaires were reviewed on the field and in the office for completeness and consistency of information. Data was entered using Statistical Package for the Social Sciences Version 22. Data was coded and cleaned before data entry. Health facilities were categorized into four namely government (secondary and tertiary) hospitals, private hospitals, primary health care centres and others (drug stores, nursing homes, traditional medicine stores). Outcome variables were client satisfaction (categorised into satisfied or dissatisfied) and quality of health care received (categorised as good or poor). The predictor variables were socio-demographic characteristics of respondents and client assessments of various aspects of services received. Association between various respondents’ characteristics and outcome variables were sought for using the Chi-Square test. Multi-variate analysis was done for factors found to be significant (p< 0.05) on bivariate analysis to identify predictors of the outcome variables. Qualitative data was analysed using ATLAS ti software version 7.^20^ The data analysis was conducted using constant comparison analyses and thematic reporting.

### Ethical considerations

The respondents were informed of the objectives of the study and its potential benefits for the health system and the state. There was no risk of harm to them. Written informed consent was obtained from each respondent prior to enrolment in the study. Ethical clearance was obtained from the Lagos State University Teaching Hospital (LASUTH) ethics committee with **Reference Number: LREC/06/10/755 (08/11/16-08/08/17)**

## Results

### Socio-demographic and socio-economic characteristics of respondents

Table 1 shows the socio-demographic and work characteristics of participants. The mean age was 37.6± 10.2 years. The largest proportion (39%) of respondents were aged 30-39 years. Over half (55%) of all respondents were females. Most of the respondents (66%) had at least a secondary school education and were married or co-habiting (77%). Almost half of the respondents were skilled workers (45%). Among the 43 FGD participants, the majority (n=33, 77%) were females, married (n=34, 79%) and Christians (n=28, 65%), and 18 (42%) had at least a secondary school education.

**Table 1.**
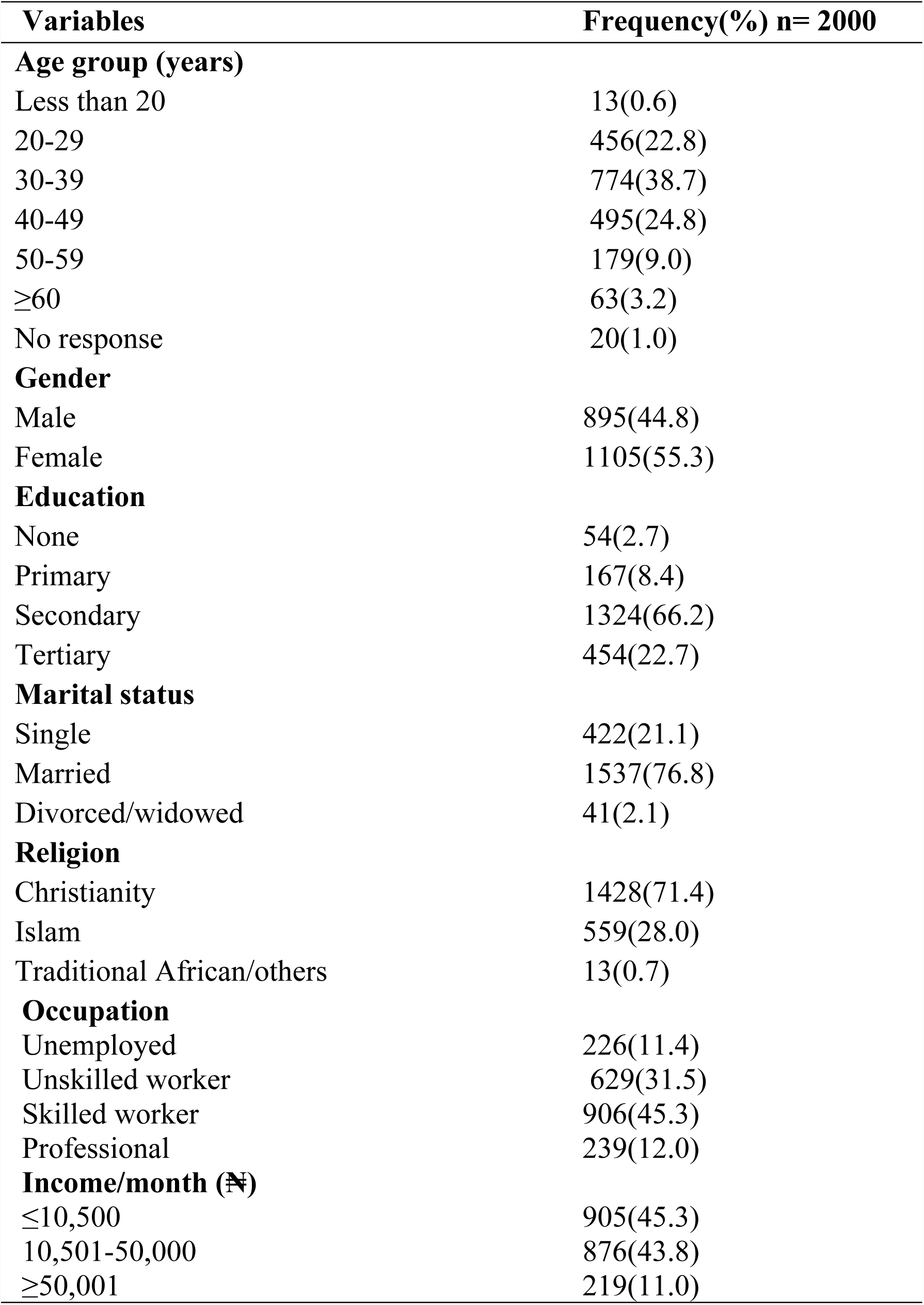
Socio-demographic and socio-economic characteristics of respondents.

### Assessment of facilities and services

Majority of respondents (98%) perceived the health facilities as being clean and considered the toilet facilities to be clean. The comfort of waiting areas in the facilities were judged to be mainly good (56%) or fair (42%). Drugs and services were considered to be cheap, (43% and 44% respectively) and 60% were of the opinion that most drugs were usually available in these facilities. Almost all the respondents (92%) expressed confidence in the skills of their health care providers. The waiting time to see the care providers was judged by over half of the respondents (60%) as being short (20 minutes or less). The majority (96%) of respondents felt the treatment received at these facilities was effective, 80% felt that the attitude of the staff was good and 95% were satisfied with the care they received (Table 2).

**Table 2.** Respondents’ assessment of quality of facilities and services.

| **Domain assessed** | **PHC Centres**<br>**N (%)** | **Secondary/tertiary facility**<br>**N (%)** | **Private Hospitals**<br>**N (%)** | **Other facilities**<br>**N (%)** | **Total**<br>**N (%)** |
| --- | --- | --- | --- | --- | --- |
| **Cleanliness** |  |  |  |  |  |
| Clean | 134 (97.8) | 790 (97.1) | 758 (99.9) | 273 (95.5) | 1995 (97.9) |
| Indifferent | 1 (0.7) | 19 (2.3) | 0 (0.0) | 12 (4.2) | 32 (1.6) |
| Dirty | 2 (1.5) | 5 (0.6) | 1 (0.1) | 1 (0.3) | 9 (0.5) |
| Total | 137(100) | 814(100) | 759(100) | 286 (100) | 1996 (100) |
| **Toilets \*** |  |  |  |  |  |
| Clean | 90 (97.8) | 563 (97.7) | 551 (100) | 72 (96.0) | 1276 (98.3) |
| Dirty | 2 (2.2) | 13 (2.3) | 0 (0.0) | 3 (4.0) | 18 (1.4) |
| Total | 92(100) | 576(100) | 551(100) | 75(100) | 1294 (100) |
| **Comfort of waiting Area** |  |  |  |  |  |
| Good | 66 (48.2) | 439 (53.9) | 471 (62.1) | 148 (51.7) | 1124 (56.3) |
| Fair | 67 (48.9) | 362 (44.5) | 285 (37.5) | 114 (39.9) | 828 (41.5) |
| Poor | 4 (2.9) | 13 (1.6) | 3 (0.4) | 24 (8.4) | 44 (2.2) |
| Total | 137(100) | 814(100) | 759(100) | 286 (100) | 1996(100) |
| **Waiting time** |  |  |  |  |  |
| Short | 61 (44.5) | 282 (34.7) | 582 (76.6) | 262 (91.9) | 1187 (59.5) |
| Average | 45 (32.8) | 302 (37.1) | 160 (21.1) | 15 (5.3) | 522 (26.2) |
| Long | 31 (22.6) | 229 (28.2) | 18 (2.4) | 8 (2.8) | 286 (14.3) |
| Total | 137 (100) | 813(100) | 760(100) | 285 (100) | 1995(100) |
| **Staff attitude** |  |  |  |  |  |
| Good | 125 (91.2) | 626 (76.9) | 684 (90.1) | 272 (95.1) | 1707 (85.5) |
| Pompous/rude | 11 (8.0) | 173 (21.2) | 72 (9.5) | 7 (2.4) | 263 (13.2) |
| Cannot assess | 1 (0.7) | 15 (1.8) | 3 (0.4) | 7 (2.4) | 26 (1.3) |
| Total | 137 (100) | 814(100) | 759(100) | 286 (100) | 1996(100) |
| **Confidence in \*\*HCP skills** |  |  |  |  |  |
| Yes | 123 (89.8) | 718 (88.2) | 725 (95.4) | 278 (97.2) | 1844 (92.3) |
| Partially | 10 (7.3) | 73 (9.0) | 23 (3.0) | 7 (2.4) | 113 (5.7) |
| No/not sure | 4 (2.9) | 23 (2.8) | 12 (1.6) | 1 (0.3) | 40 (2.0) |
| Total | 137 (100) | 814(100) | 760(100) | 286 (100) | 1997(100) |
| **Effective Treatment** |  |  |  |  |  |
| Yes | 130 (94.9) | 775 (95.3) | 739 (97.5) | 277 (97.2) | 1921 (96.4) |
| No | 3 (2.2) | 24 (3.0) | 14 (1.8) | 3 (1.1) | 44 (2.2) |
| Not sure | 4 (2.9) | 14 (1.7) | 5 (0.7) | 5 (1.8) | 28 (1.4) |
| Total | 137 (100) | 813(100) | 758(100) | 285 (100) | 1993(100) |
| **Availability of drugs** |  |  |  |  |  |
| All | 23 (16.8) | 93 (11.4) | 241 (31.8) | 146 (51.0) | 503 (25.2) |
| Most | 83 (60.6) | 542 (66.7) | 465 (61.3) | 99 (34.6) | 1189 (59.6) |
| Few/None | 31 (22.6) | 178 (21.9) | 53 (6.9) | 41 (14.3) | 303 (15.2) |
| Total | 137 (100) | 813(100) | 759 (100) | 286 (100) | 1995(100) |
| **Cost of card** | **(₦)** |  |  |  |  |
| None | 49 (35.8) | 119 (14.7) | 37 (4.9) | 256 (89.8) | 461 (23.2) |
| <500 | 54 (39.4) | 425 (52.4) | 188 (25.0) | 12 (4.3) | 679 (34.1) |
| ≥500 | 34 (24.8) | 268 (33.0) | 528 (70.1) | 17 (6.0) | 847 (42.7) |
| Total | 137 (100) | 812 (100) | 753(100) | 285 (100) | 1987(100) |
| **Cost of Drugs** |  |  |  |  |  |
| Cheap | 75 (54.7) | 379 (46.7) | 187 (24.6) | 219 (76.6) | 860 (43.1) |
| Fair | 42 (30.7) | 321 (39.5) | 332 (43.7) | 48 (16.8) | 743 (37.3) |
| Expensive | 20 (14.6) | 112 (13.8) | 240 (31.6) | 19 (6.6) | 391 (19.6) |
| Total | 137(100) | 812 (100) | 759(100) | 286(100) | 1994(100) |
| **Cost of Services** |  |  |  |  |  |
| Cheap | 83 (60.6) | 366 (45.0) | 225 (29.6) | 213 (74.5) | 887 (44.4) |
| Fair | 41 (29.9) | 387 (47.5) | 389 (51.3) | 57 (19.9) | 874 (43.8) |
| Expensive | 13 (9.5) | 61 (7.5) | 145 (19.1) | 16 (5.6) | 235 (11.8) |
| Total | 137 (100) | 814 (100) | 759(100) | 286 (100) | 1996(100) |
| **Satisfaction** |  |  |  |  |  |
| Satisfied | 127 (92.7) | 743 (91.3) | 741 (97.6) | 280 (97.9) | 1891 (94.7) |
| Dissatisfied | 10 (7.3) | 71 ( 8.7) | 18 (2.4) | 6 (2.1) | 105 (5.3)) |
| Total | 137 (100) | 814(100) | 759(100) | 286 (100) | 1996(100) |

\*Only health facilities that had toilets
\*\*HCP = Health Care Provider

The FGD participants generally perceived that private-owned hospitals were more conducive, more attractive and cleaner compared to government owned hospitals in several aspects (the environment, waiting areas, toilets and consultation rooms).

A participant explained that clients sometimes slept comfortably outside the private hospital he visits because the environment is conducive and neat.

> *“…it is ok (the environment), they perform Caeserean sections there, when you get there, you will meet all of them [patients] outside receiving fresh air, even most of those who came to visit the patients, mostly sleep outside. If the place is not ok, they won’t sleep there.” –* ***Badagry_female_no education_58years_married***
>
> *Another female participant in Ojo attested to the cleanliness of the facility she used “…everything is also ok, all their toilets are ok, and the doctor’s consulting room as well is ok”.*

*In contrast, p*articipants from Badagry complained about the government owned health facility within their location. They complained that the environment was not clean, the toilets were very dirty and the grass in the environment around the toilet was overgrown. A female participant from Badagry explained that the government-owned hospital was always infested with mosquitoes, because the environment was not well kept, and the mosquito nets had not been replaced.

> She stated, “*…secondly mosquitoes, I can’t sleep, and they said there is net, you will just see some nets, some are already torn. They won’t replace … …. You can’t sleep, mosquitoes will continue to bite you, and the fever will get worse”(****Badagry_female_secondary_45years_married).***

As regards public/government owned health facilities, more participants mentioned that the modalities of operations were stressful. The process of walking around from one point to the other was mentioned as one of the most stressful things patients had to undergo during a clinic visit. A participant said:

> *“…it’s not easy at all in government hospitals. When you first arrive, you might need to obtain a card from one of the rooms and you will have to queue. The place where you are to receive drugs at the pharmacy, might be located in the second or the third block. If they now discover that it is operation that you want to do, they might now tell you to go and take the X-ray or some other test, before you now move from that point to the section where the operation will be performed. This really stresses some people. If it is a private facility, all that you need to do will be taken care off in one place. That is why that one is different, but for government hospital, the distance you will cover there, it’s not small at all, the distance you will cover there [within the hospital premises] will be similar to taking public transport, and for someone who is sick and is not feeling well and going through pain and distress and who is in God’s hands…. The departments are always too far apart. You go to one place, and when you finish there, you have to go to another place to pay and then another place.” –* ***Ikeja_male_secondary_38years_single***

A participant’s opinion of private health facilities is shown below.

> *“…At the private hospital that I use, at Badagry, for anything you need, it is the nurse that will get it for you, but the government their problem is too much, the units are too far apart.”“… in private hospital, units/departments are not far flung, but in the general hospital, if the reception is here, the doctor’s office is somewhere else, you will receive your drugs in another place, … … … it [the general hospital] is very big.*
>
> ***-Ikeja_male_tertiary_married***

Another respondent opined as follows

> *“: You will walk around [in government hospitals], but in private, you will just relax, they will ask you everything they want to ask you, if they want to treat you, they will just go to their pharmacy, the nurses will do that, you don’t need to walk around, you will just stay in one place, they [the staff] will do what they want to do.” -* ***Badagry_female_secondary_20 years_single***

### Determinants of client satisfaction

Almost 95% of the respondents were satisfied with the services they received. Satisfaction with services at the health facilities was significantly associated with respondents’ marital status, occupation, income and the type of facility patronised as usual source of care (p<0.05). There was no association between satisfaction and respondents’ age group, gender, educational level and religion. (Table 3). Table 4 shows the association between respondents’ perception of service characteristics and client satisfaction. Cost of services, cleanliness of the facility, short waiting time and positive staff attitudes were significantly associated with client satisfaction. The predictive factors of client satisfaction on logistic regression, the were the use of government owned hospitals (OR 5.78, 95% confidence intervals, 2.433 −13.700, p < 0.001) and PHCs (OR 5.0 95% confidence intervals, 1.715-14.286, p=0.003). Other predictor variables were waiting time (short/average), costs of drugs, cost of services and staff attitudes and confidence in the health care provider (Table 5).

**Table 3.** Association between respondents’ socio-demographic characteristics and client satisfaction.

| **Socio-demographic variable** | **Satisfied** | **n (%)** | **Dissatisfied** | **Total** | **Test of significance** |
| --- | --- | --- | --- | --- | --- |
|  |  |  | **n (%)** | **n (%)** |  |
| **Age group (years)** |  |  |  |  |  |
| <20 | 11 (84.6) |  | 2 (15.4) | 13 (100) | X<sup>2</sup>=10.77<br>df=5<br>p=0.056 |
| 20-29 | 422 (92.5) |  | 34 (7.5) | 456 (100) |  |
| 30-39 | 740 (95.7) |  | 33 (4.3) | 773 (100) |  |
| 40-49 | 467 (94.3) |  | 28 (5.7) | 495 (100) |  |
| 50-59 | 174 (97.2) |  | 5 (2.8) | 179 (100) |  |
| ≥ 60 | 61 (96.8) |  | 2 (3.2) | 63 (100) |  |
| Total | 1875 (94.7) |  | 104 (5.3) | 1979 (100) |  |
| **Gender** |  |  |  |  |  |
| Male | 842 (94.1) |  | 53 (5.9) | 895 (100) | X<sup>2</sup>=1.458<br>df=1<br>p=0.227 |
| Female | 1052 (95.3) |  | 52 (4.7) | 1104 (100) |  |
| Total | 1894 (94.7) |  | 105 (5.3) | 1999 (100) |  |
| **Education** |  |  |  |  |  |
| No formal | 52 (96.3) |  | 2 (3.7) | 54 (100) | X<sup>2</sup>=4.089<br>df=3<br>p=0.252 |
| Primary | 160 (95.8) |  | 7 (4.2) | 167 (100) |  |
| Secondary | 1260 (95.2) |  | 64 (4.8) | 1324 (100) |  |
| Tertiary | 421 (92.9) |  | 32 (7.1) | 453 (100) |  |
| Total | 1893 (94.7) |  | 105 (5.3) | 1998 (100) |  |
| **Marital status** |  |  |  |  |  |
| Single | 386 (91.5) |  | 36 (8.5) | 422 (100) | X<sup>2</sup>=12.283<br>df=2<br>**p=0.002** |
| Married | 1470 (95.7) |  | 66 (4.3) | 1536 (100) |  |
| Divorced/widowed | 38 (92.7) |  | 3 (7.3) | 41 (100) |  |
| Total | 1894 (94.7) |  | 105 (5.3) | 1999 (100) |  |
| **Religion** |  |  |  |  |  |
| Christianity | 1352 (94.7) |  | 75 (5.3) | 1427 (100) | X<sup>2</sup>=0.00<br>df=1<br>p=1.0 |
| Islam/others | 541 (94.7) |  | 30 (5.3) | 571 (100) |  |
| Total | 1893 (94.7) |  | 105 (5.3) | 1998 (100) |  |
| **Occupation** |  |  |  |  |  |
| Unemployed | 200 (90.1) |  | 22 (9.9) | 222 (100) | X<sup>2</sup>=10.94<br>df=3<br>P=0.012 |
| Unskilled work | 599 (95.2) |  | 30 (4.8) | 629 (100) |  |
| Skilled work | 865 (95.5) |  | 41 (4.5) | 906 (100) |  |
| Professionals | 226 (95.0) |  | 12 (5.0) | 238 (100) |  |
| Total | 1890 (94.7) |  | 105 (5.3) | 1995 (100.0) |  |
| **Income (N)** |  |  |  |  |  |
| ≤10,500 | 856 (94.6) |  | 49 (5.4) | 905 (100) | X<sup>2</sup>= 12.766<br>df= 5<br>**p=0.026** |
| 10,501-50,000 | 824 (90.4) |  | 51 (9.6) | 875 (100) |  |
| >50,000 | 212 (97.6) |  | 5 (2.4) | 219 (100) |  |
| Total | 1894 (94.7) |  | 105 (5.3) | 1999 (100.0) |  |
| **Sources of health care services** |  |  |  |  |  |
| Government hospitals | 743 (91.3) |  | 71 (8.7) | 814 (100) | X<sup>2</sup>= 39.167<br>df= 3<br>**p< 0.001** |
| Private hospitals | 741 (97.6) |  | 18 (2.4) | 759 (100) |  |
| Primary health centre | 127 (92.7) |  | 10 (7.3) | 137 (100) |  |
| Others | 280 (97.9) |  | 6 (2.1) | 286 (100) |  |
| Total | 1891 (94.7) |  | 105 (5.3) | 1996 (100.0) |  |

**Table 4.**
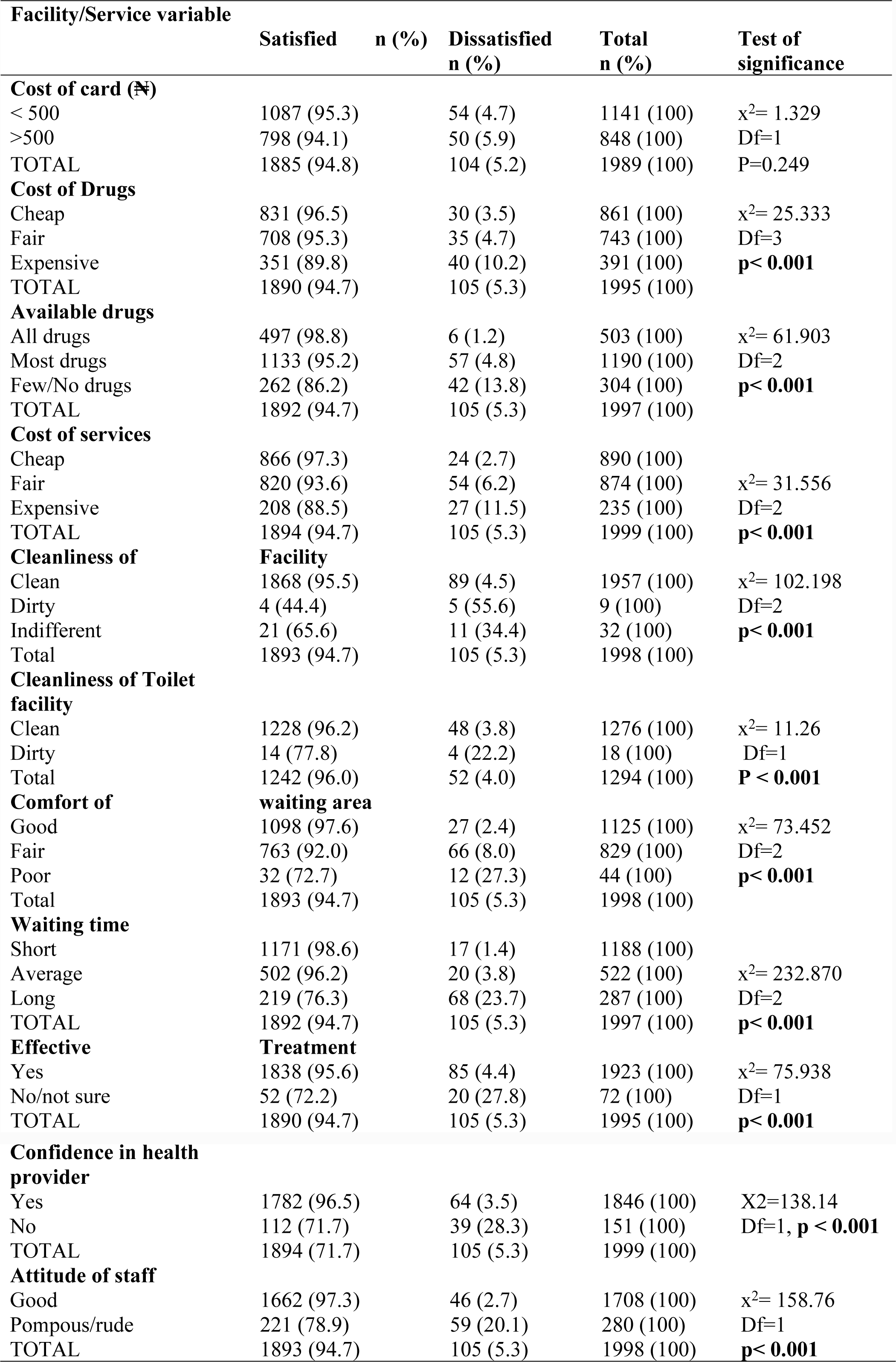
Association between service characteristics and client satisfaction.

| Facility/Service variable | Satisfied | n (%) | Dissatisfied<br>n (%) | Total<br>n (%) | Test of<br>significance |
| --- | --- | --- | --- | --- | --- |
| **Cost of card (₦)** |  |  |  |  |  |
| < 500 | 1087 (95.3) | | 54 (4.7) | 1141 (100) | $\chi^2= 1.329$ |
| >500 | 798 (94.1) |  | 50 (5.9) | 848 (100) | Df=1 |
| TOTAL | 1885 (94.8) |  | 104 (5.2) | 1989 (100) | P=0.249 |
| **Cost of Drugs** |  |  |  |  |  |
| Cheap | 831 (96.5) | | 30 (3.5) | 861 (100) | $\chi^2= 25.333$ |
| Fair | 708 (95.3) |  | 35 (4.7) | 743 (100) | Df=3 |
| Expensive | 351 (89.8) |  | 40 (10.2) | 391 (100) | **p< 0.001** |
| TOTAL | 1890 (94.7) |  | 105 (5.3) | 1995 (100) |  |
| **Available drugs** |  |  |  |  |  |
| All drugs | 497 (98.8) | | 6 (1.2) | 503 (100) | $\chi^2= 61.903$ |
| Most drugs | 1133 (95.2) |  | 57 (4.8) | 1190 (100) | Df=2 |
| Few/No drugs | 262 (86.2) |  | 42 (13.8) | 304 (100) | **p< 0.001** |
| TOTAL | 1892 (94.7) |  | 105 (5.3) | 1997 (100) |  |
| **Cost of services** |  |  |  |  |  |
| Cheap | 866 (97.3) |  | 24 (2.7) | 890 (100) |  |
| Fair | 820 (93.6) | | 54 (6.2) | 874 (100) | $\chi^2= 31.556$ |
| Expensive | 208 (88.5) |  | 27 (11.5) | 235 (100) | Df=2 |
| TOTAL | 1894 (94.7) |  | 105 (5.3) | 1999 (100) | **p< 0.001** |
| **Cleanliness of Facility** |  |  |  |  |  |
| Clean | 1868 (95.5) | | 89 (4.5) | 1957 (100) | $\chi^2= 102.198$ |
| Dirty | 4 (44.4) |  | 5 (55.6) | 9 (100) | Df=2 |
| Indifferent | 21 (65.6) |  | 11 (34.4) | 32 (100) | **p< 0.001** |
| Total | 1893 (94.7) |  | 105 (5.3) | 1998 (100) |  |
| **Cleanliness of Toilet facility** |  |  |  |  |  |
| Clean | 1228 (96.2) | | 48 (3.8) | 1276 (100) | $\chi^2= 11.26$ |
| Dirty | 14 (77.8) |  | 4 (22.2) | 18 (100) | Df=1 |
| Total | 1242 (96.0) |  | 52 (4.0) | 1294 (100) | **P < 0.001** |
| **Comfort of waiting area** |  |  |  |  |  |
| Good | 1098 (97.6) | | 27 (2.4) | 1125 (100) | $\chi^2= 73.452$ |
| Fair | 763 (92.0) |  | 66 (8.0) | 829 (100) | Df=2 |
| Poor | 32 (72.7) |  | 12 (27.3) | 44 (100) | **p< 0.001** |
| Total | 1893 (94.7) |  | 105 (5.3) | 1998 (100) |  |
| **Waiting time** |  |  |  |  |  |
| Short | 1171 (98.6) |  | 17 (1.4) | 1188 (100) |  |
| Average | 502 (96.2) | | 20 (3.8) | 522 (100) | $\chi^2= 232.870$ |
| Long | 219 (76.3) |  | 68 (23.7) | 287 (100) | Df=2 |
| TOTAL | 1892 (94.7) |  | 105 (5.3) | 1997 (100) | **p< 0.001** |
| **Effective Treatment** |  |  |  |  |  |
| Yes | 1838 (95.6) | | 85 (4.4) | 1923 (100) | $\chi^2= 75.938$ |
| No/not sure | 52 (72.2) |  | 20 (27.8) | 72 (100) | Df=1 |
| TOTAL | 1890 (94.7) |  | 105 (5.3) | 1995 (100) | **p< 0.001** |

**Confidence in health provider**
|  |  |  |  |  |
| --- | --- | --- | --- | --- |
| Yes | 1782 (96.5) | 64 (3.5) | 1846 (100) | X<sup>2</sup>=138.14 |
| No | 112 (71.7) | 39 (28.3) | 151 (100) | Df=1, **p < 0.001** |
| TOTAL | 1894 (71.7) | 105 (5.3) | 1999 (100) |  |

**Attitude of staff**
|  |  |  |  |  |
| --- | --- | --- | --- | --- |
| Good | 1662 (97.3) | 46 (2.7) | 1708 (100) | x<sup>2</sup>= 158.76 |
| Pompous/rude | 221 (78.9) | 59 (20.1) | 280 (100) | Df=1 |
| TOTAL | 1893 (94.7) | 105 (5.3) | 1998 (100) | **p< 0.001** |

**Table 5.**
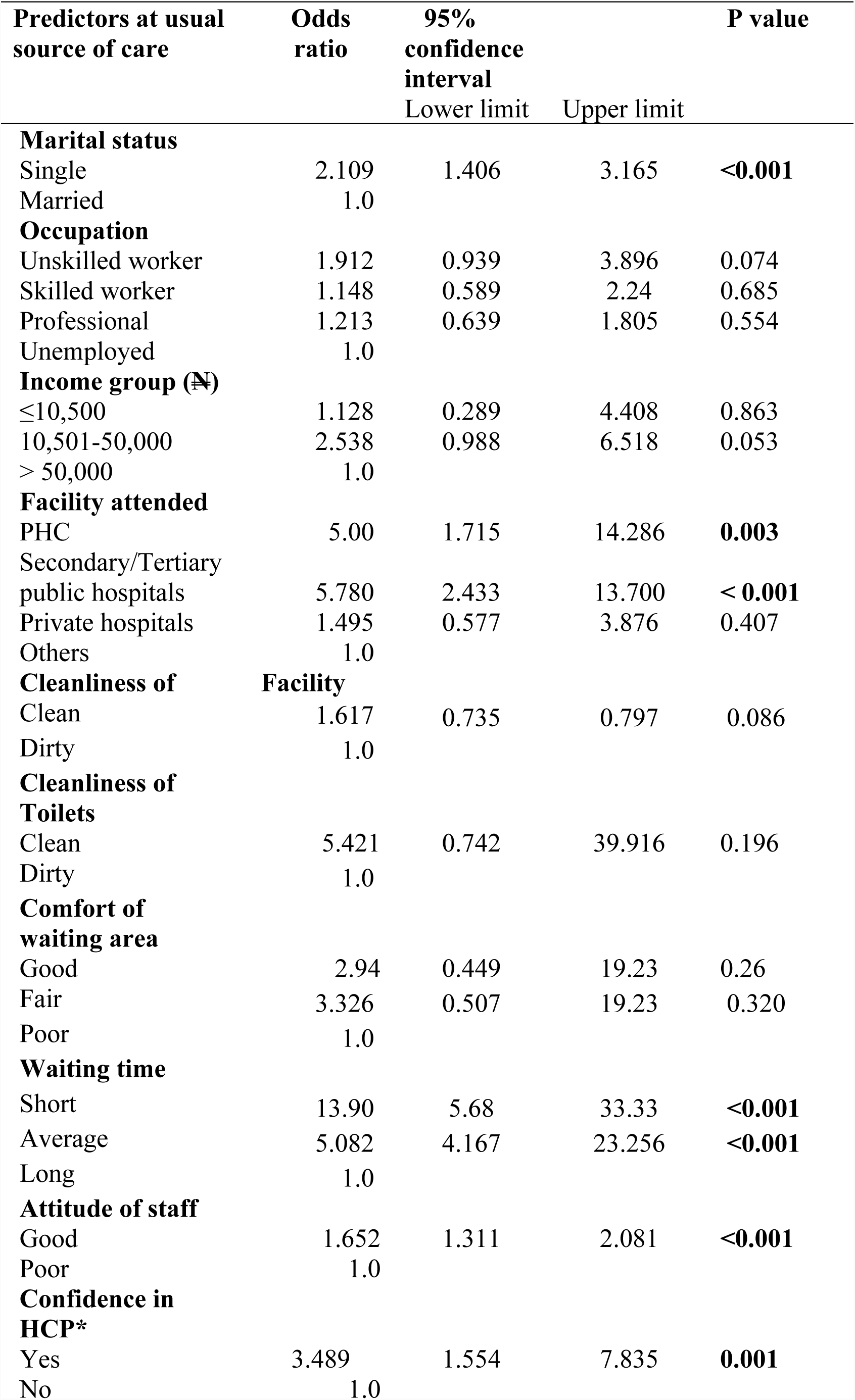
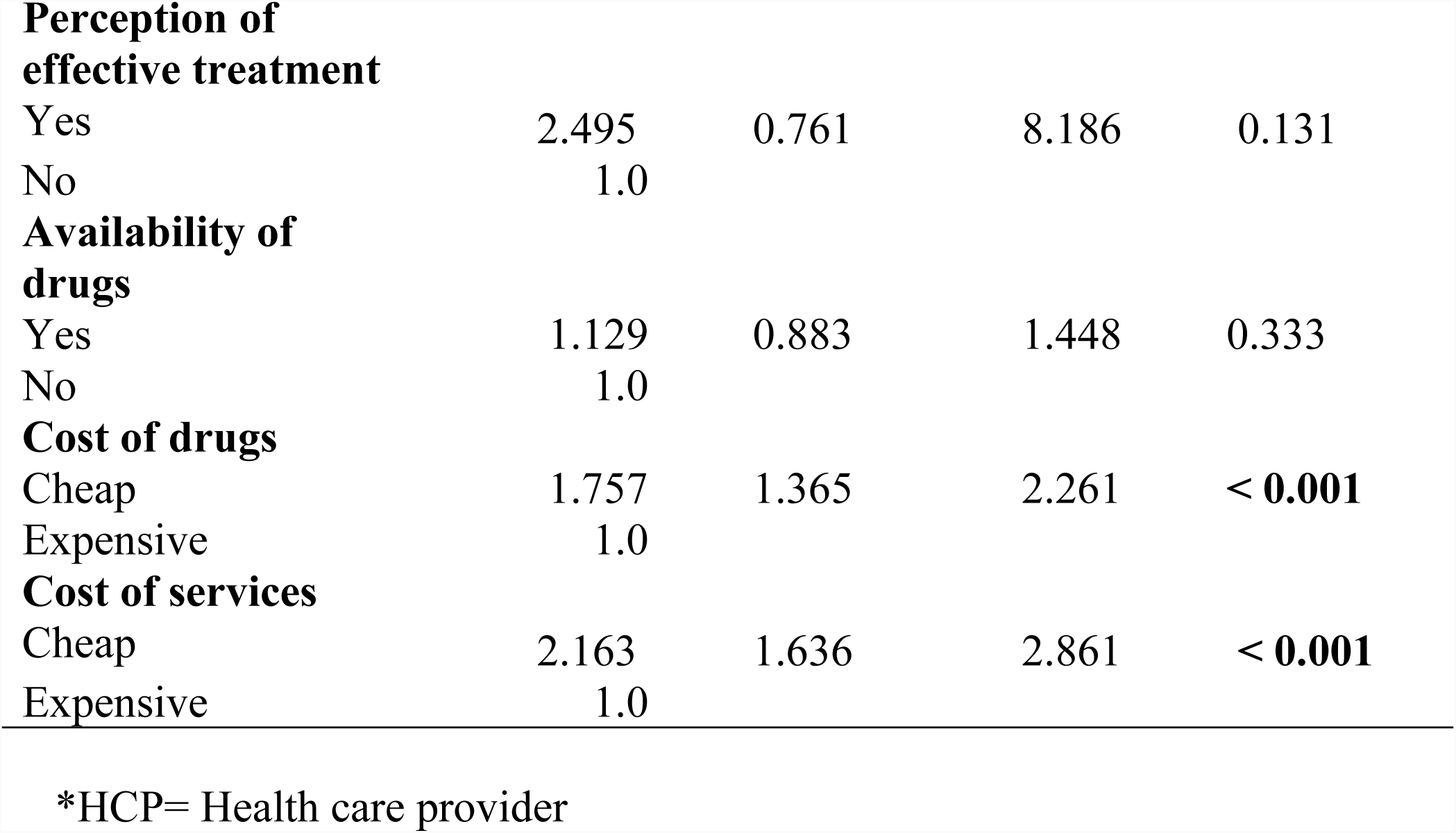
Predictors of client satisfaction.

### Determinants of quality of care

About 85% of the respondents perceived the quality of services they received to be of good quality. Respondents’ characteristics found to be significantly associated with perception of good service quality were gender, income and the source of health care but age, and marital status were not (Table 6). All the service components assessed except staff attitudes and costs of services (card, drugs and all services combined) were found to be significantly associated with perception of good service quality (Table 7). Male gender and participants who earned low income were found to be predictors of poor quality. Factors found to be predictors of good quality were use of higher level of health care, confidence in the health care providers, perceived effectiveness of treatment received, comfort of waiting area and availability of drugs (Table 8). A significant correlation (Spearman’s correlation, r=0.145, p=0.001) was found between client satisfaction and service quality.

**Table 6.**
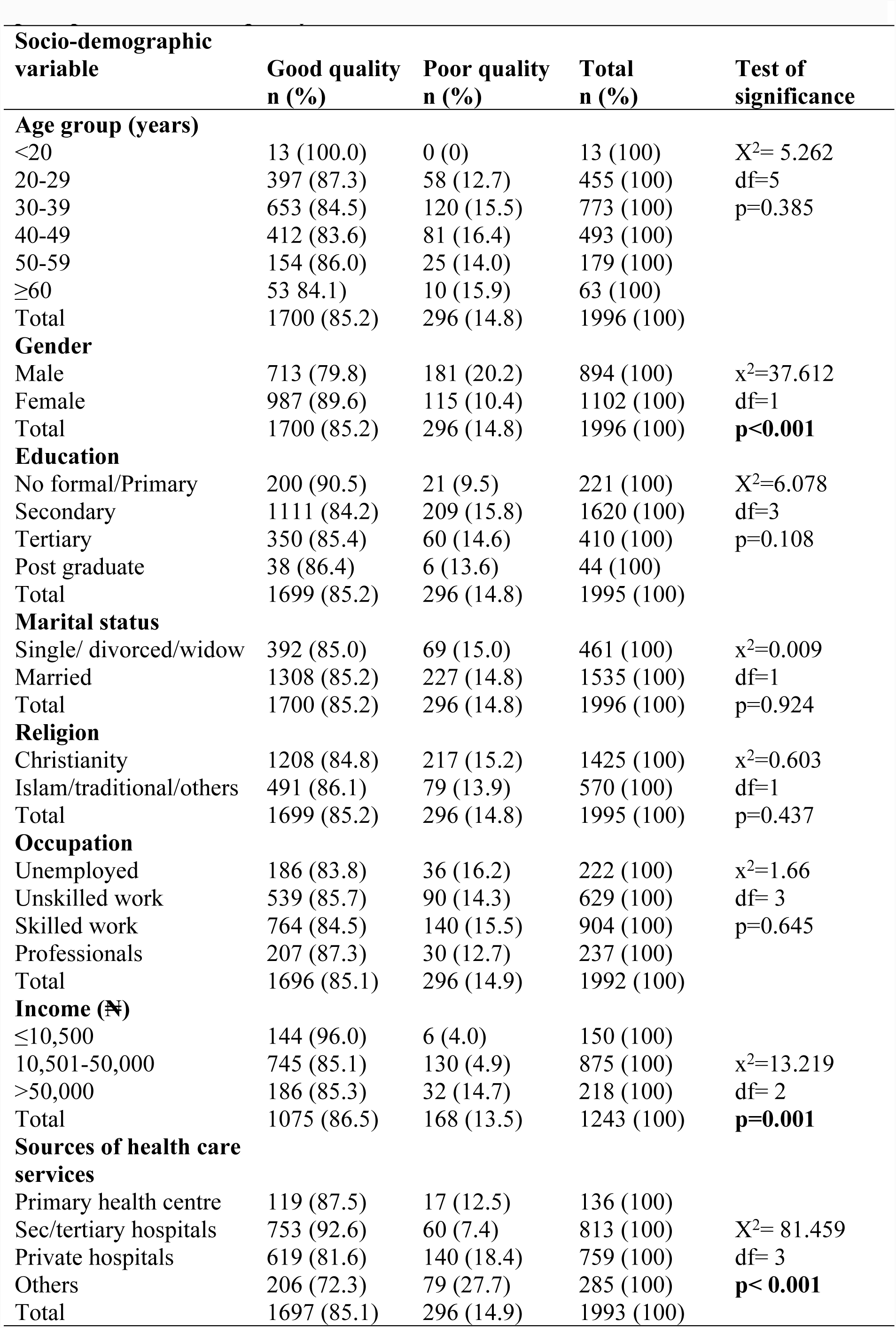
Association between respondents’ socio-demographic characteristics and perception of service quality.

**Table 7.** Association between service characteristics respondents’ perception of service quality.

| Variable | Good quality n(%) | Poor quality n(%) | Total n (%) | Significance |
| --- | --- | --- | --- | --- |
| **Cost of card (₦)** |  |  |  |  |
| < 500 | 978 (85.9) | 161 (14.1) | 1139 (100) | $\chi^2= 0.60$ |
| >500 | 715 (84.5) | 131 (15.5) | 846 (100) | Df=1 |
| TOTAL | 1693 (85.3) | 283 (14.7) | 1985 (100) | P =0.44 |
| **Cost of Drugs** |  |  |  |  |
| Cheap | 723 (84.1) | 137 (15.9) | 860 (100) | $\chi^2= 3.706$ |
| Fair | 646 (87.2) | 95 (12.8) | 741 (100) | Df=2 |
| Expensive | 328 (83.9) | 62 (16.1) | 391 (100) | P=0.157 |
| TOTAL | 1697 (85.2) | 295 (14.8) | 1992 (100) |  |
| **Available drugs** |  |  |  |  |
| All drugs | 410 (81.7) | 92 (18.3) | 502 (100) | $\chi^2= 14.655$ |
| Most drugs | 1042 (87.7) | 146 (12.3) | 1188 (100) | Df=2 |
| Few/No drugs | 247 (81.2) | 57 (18.8) | 304 (100) | **P=0.001** |
| TOTAL | 1699 (85.2) | 295 (14.8) | 1994 (100) |  |
| **Cost of services** |  |  |  |  |
| Cheap | 761 (85.7) | 127 (14.3) | 888 (100) |  |
| Fair | 751 (86.0) | 122 (14.0) | 873 (100) | $\chi^2= 5.816$ |
| Expensive | 187 (79.9) | 47 (20.1) | 234 (100) | Df=2 |
| TOTAL | 1699 (85.2) | 296 (14.2) | 1995 (100) | P=0.055 |
| **Cleanliness of Facility** |  |  |  |  |
| Clean | 1672 (85.6) | 282 (14.4) | 1954 (100) | $\chi^2= 10.841$ |
| Dirty | 27 (65.9) | 14 (34.1) | 41 (100) | Df=1 |
| Total | 1699 (85.2) | 296 (14.8) | 1995 (100) | **P=0.001** |
| **Toilet facility** |  |  |  |  |
| Clean | 1113 (87.4) | 161 (12.6) | 1274 (100) | Fisher's exact |
| Dirty | 12 (66.7) | 6 (33.3) | 18 (100) |  |
| Total | 1125 (87.1) | 167 (12.9) | 1292 (100) | **p=0.021** |
| **Comfort of waiting area** |  |  |  |  |
| Good | 984 (87.6) | 139 (12.4) | 1123 (100) | $\chi^2= 17.185$ |
| Fair | 684 (82.6) | 144 (17.4) | 828 (100) | Df=2 |
| Poor | 31 (70.5) | 13 (29.5) | 44 (100) | **P < 0.001** |
| Total | 1699 (85.2) | 296 (14.8) | 1995 (100) |  |
| **Waiting time** |  |  |  |  |
| Short | 984 (83.0) | 202 (17.0) | 1186 (100) |  |
| Average | 474 (91.3) | 45 (8.7) | 519 (100) | $\chi^2= 20.584$ |
| Long | 241 (84.0) | 46 (16.0) | 287 (100) | Df=2 |
| TOTAL | 1699 (85.3) | 293 (14.7) | 1992 (100) | **p< 0.001** |
| **Effective Treatment** |  |  |  |  |
| Yes | 1647 (85.8) | 273 (14.2) | 1920 (100) | $\chi^2= 13.415$ |
| No | 50 (69.4) | 22 (30.6) | 72 (100) | Df=1 |
| TOTAL | 1697 (85.2) | 295 (14.5) | 1992 (100) | **p< 0.001** |
| **Confidence in health provider** |  |  |  |  |
| Yes | 1593 (86.4) | 250 (13.6) | 1843 (100) | $\chi^2= 13.584$ |
| No | 107 (74.8) | 36 (25.2) | 143 (100) | Df= 1 |
| TOTAL | 1700 (85.6) | 286 (14.4) | 1986 (100) | **p< 0.001** |
| **Attitude of staff** | | | | $\chi^2 = 2.292$ |
| Good | 1461 (85.7) | 244 (14.3) | 1705 (100) | Df=1 |
| Pompous/rude | 238 (82.1) | 52 (17.9) | 290 (100) | P= 0.130 |
| TOTAL | 1699 (85.1) | 296 (14.9) | 1995 (100) |  |

**Table 8.** Predictors of Respondents assessment of service quality.

| Variable | Odds ratio | 95% confidence interval |  | P value |
| --- | --- | --- | --- | --- |
|  |  | Lower limit | Upper limit |  |
| **Gender** |  |  |  |  |
| Male | 0.468 | 0.364 | 0.602 | **<0.001** |
| Female | 1.0 |  |  |  |
| **Income (₦)** |  |  |  |  |
| ≤10,500 | 0.363 | 0.141 | 0.934 | **0.036** |
| 10,501-50,000 | 1.040 | 0.648 | 1.669 | 0.871 |
| >50,000 | 1.0 |  |  |  |
| **Facility used** |  |  |  |  |
| PHC Centre | 1.867 | 1.066 | 3.269 | **< 0.001** |
| Secondary/tertiary government hospitals | 24.689 | 2.207 | 276.147 | **0.009** |
| Private hospitals | 4.629 | 3.202 | 6.692 | **< 0.001** |
| Others | 1.0 |  |  |  |
| **Clean facility** |  |  |  |  |
| Clean | 1.218 | 0.942 | 1.575 | 0.132 |
| Dirty | 1.0 |  |  |  |
| **Cleanliness of Toilets** |  |  |  |  |
| Yes | 1.015 | 0.909 | 1.134 | 0.789 |
| No | 1.0 |  |  |  |
| **Waiting time** |  |  |  |  |
| Short/average | 1.120 | 0.800 | 1.572 | 0.506 |
| Long | 1.0 |  |  |  |
| **Comfort of waiting area** |  |  |  |  |
| Good | 2.817 | 1.440 | 5.490 | **0.002** |
| Fair | 1.916 | 0.981 | 3.740 | 0.057 |
| Poor | 1.0 |  |  |  |
| **Confidence in Health provider** |  |  |  |  |
| Yes | 2.234 | 1.509 | 3.308 | **<0.001** |
| No | 1.0 |  |  |  |
| **Effectiveness of treatment** |  |  |  |  |
| Yes | 1.835 | 1.060 | 3.179 | **0.030** |
| No | 1.0 |  |  |  |
| **Availability of drugs** |  |  |  |  |
| Yes | 1.120 | 1.007 | 1.244 | **0.036** |
| No | 1.0 |  |  |  |

\*\*HCP = Health Care Provider

## Discussion

This study investigated client satisfaction and quality of care received amongst residents of Lagos State, Nigeria. These factors have a great potential on service utilization. A qualitative study from Uganda reported high costs, poor attitude of staff, non-availability of services as barriers to utilization of services and there was the perception that public health faculties offered low quality care.^21^ Besides, a Nigerian study had shown that utilization of health services was higher when the perceived quality was good.^11^ This is important in Nigeria and other countries where health services are “essentially consumer goods” paid for largely through out-of-pocket mechanisms and as such clients should therefore get maximum value for money spent. Furthermore, the quality of service may in part determine the choice between orthodox and non-orthodox providers. Nine out of ten respondents were satisfied with the services received and 85% perceived the services was of good quality. The significant correlation (r=0.145, p =0.01) between both outcomes further indicate that both should be addressed simultaneously in the provisions of health services. In addition, the predictor variables of respondents and service characteristics which showed significant associations with the outcome variables were similar thus strengthening the evidence from the study.

The proportion (95%) of clients who were satisfied in this study was far higher than the 55% reported from Ile-Ife.^12^ Being single was predictive of client satisfaction but we did not find other predictive factors such as education as reported from Ile-Ife or age and gender as reported from Ilorin.^16^ While the reasons for such non-concordance are not known to the authors they may have to do in part with differences in the study populations. Furthermore, the inability to identify many respondents’ characteristics to be predictive of client satisfaction is supported by the work of Tucker et al^7^ who had reported that clients’ socio-demographic characteristics account for less than 1% of the variance of client satisfaction. It may also be that identification of the factors require more rigorous methods beyond the scope of the present study. Our study showed that service characteristics such as cost of service, costs of drugs, staff attitudes and confidence in the health care providers were predictive of client satisfaction in consonance with the works of other researchers.^2.3,7^ These findings show that client satisfaction is achievable if adequate attention is paid to delivering good and affordable services.

The level of good service quality(85%) in this study was higher than values reported from Nnewi (65%),^22^ Ile-Ife (72%),^12^ but similar to Lagos (88%)^16^ and Bangladesh (90%).^23^ The proportion of respondents in Bangladesh who rated the facilities to be clean was 72%; adequacy of time spent with physicians (54%) and opportunity to ask questions (84%). Moreover, the proportion of respondents who rated the environment clean in this study was much higher than the 46% reported from Benin city.^24^ Comfort of the waiting area, effectiveness of treatment, availability of drugs and confidence in the health care provider were found to be predictive of good service quality. The study did not find use of private facilities to be predictive of client satisfaction, which is similar to a report from Abeokuta, Nigeria^25^ and in contrast to the views expressed by the FGD participants. The factors found to be associated with service quality in that study^25^ are similar to our findings.

Using the SERVIQUAL model domains,^8^ this study found that four of these were rated very highly; tangibles (environment), responsiveness (promptness of service), assurance (explanation of health conditions and knowledge) and reliability (competence) in concordance with a study at the out-patient clinics of a general hospital in Lagos.^16^ It is to be noted that patients in diverse health facility settings report differently their expectations on the importance of domains of quality. In teaching hospitals in south west Nigeria, reliability dimension was the most important^14^ whereas at general hospitals in the same region, empathy was the most important.^15^ This may be related in part to the more severe illnesses presenting at teaching hospitals and staff attitudes. Using the Donabedian model,^1^ it can be said that the “structure and process dimensions” of health services offered in Lagos State were good. The cross-sectional nature of the study did not allow for assessment of the “outcome dimensions”.

What factors were responsible for the favourable assessment by the respondents? First the environments of the facilities were found to be clean and comfortable including the toilets. The environment is the first contact of the client with the health facility. The second factor lies in a number of service characteristics for example short waiting time, affordable fees and availability of drugs. These are likely to meet client expectations and such clients will not only return but perhaps refer others.

### Limitations of the study

The study limitations included social desirability bias as respondents are known to speak positively to interviewers. Careful explanation of the objectives and the anonymity required helped to minimize this. In addition, recall bias is a known limitation of questionnaire-based surveys.

### Strengths of the study

The study has several strengths. First, the sample size is large and robust allowing for valid inferences about the study outcomes to be made. Being a community-based study enables the study to investigate the key issues and include clients who have and those who have not used health facilities unlike many others which are hospital-based. In addition, the study design included users of private and public health facilities and across multiple levels of care.

## Conclusion

The respondents rated several aspects of health services very highly. Ninety-five percent of the respondents were satisfied with the services received and 85% rated the services to be of good quality. We recommend that the adequate attention should continue to be paid to ensure that the environments of health facilities are clean and comfortable for clients. In addition, the management of government health facilities should address the issues of prolonged waiting time to improve client satisfaction and experiences. This can be achieved using staggered appointments and two-way referral system. In addition, health workers undergo serial retraining on communication skills and inter-personal relationship to improve service quality. Further research to identify more factors affecting client satisfaction and service quality is recommended.

## Acknowledgements

The authors wish to acknowledge the help rendered by Dr Temitope Durojaiye, Dr Anifowose, research assistants and the members of the public who were kind enough to be part of the study.

## Conflict of interest

The authors declare there are no conflicts of interest.

### Funding

The study was funded by the Tertiary Education Trust Fund

### Authors’ contributions

AMR, OEO, supervised data collection, performed data analysis and wrote the manuscript OOO initiated the concept of the study, was involved in data analysis, wrote the manuscript and provided overall direction for the study.

All authors approved the manuscript.

